# Sparse Bayesian Learning for Predicting Phenotypes and Ranking Influential Markers in Yeast

**DOI:** 10.1101/489245

**Authors:** Maryam Ayat, Michael Domaratzki

## Abstract

Genomic selection and genome-wide association studies are two related problems that can be applied to the plant breeding industry. Genomic selection is a method to predict phenotypes (i.e., traits) such as yield and drought resistance in crops from high-density markers positioned throughout the genome of the varieties. In this paper, we employ employ sparse Bayesian learning as a technique for genomic selection and ranking markers based on their relevance to a trait, which can aid in genome-wide association studies. We define and explore two different forms of the sparse Bayesian learning for predicting phenotypes and identifying the most influential markers of a trait, respectively. In particular, we introduce a new framework based on sparse Bayesian and ensemble learning for ranking influential markers of a trait. Then, we apply our methods on a real-world *Saccharomyces cerevisiae* dataset, and analyse our results with respect to existing related works, trait heritability, as well as the accuracies obtained from the use of different kernel functions including linear, Gaussian, and string kernels. We find that sparse Bayesian methods are not only as good as other machine learning methods in predicting yeast growth in different environments, but are also capable of identifying the most important markers, including both positive and negative effects on the growth, from which biologists can get insight. This attribute can make our proposed ensemble of sparse Bayesian learners favourable in ranking markers based on their relevance to a trait.

## Introduction

Genomic Selection (GS) and Genome-Wide Association Study (GWAS) are two related problems in bioinformatics that represent two different aspects of a quantitative trait. GS predicts phenotypes such as growth and fertility in livestocks [1, 2], and yield and drought resistance in crops [3], using genetic information of individuals, that is, sequences of genome-wide molecular markers. Single Nucleotide Polymorphisms (SNPs) are the most common type of genetic markers. GS is ideal for complex traits, which are controlled by many genes with different effects across the genome [4]. GS in plants or animals are mainly used in the breeding industry to facilitate the selection of superior genotypes and accelerate the breeding cycle [5, 6].

On the other hand, GWAS helps to investigate the genetic architecture of causal interpretation of phenotypic variations in humans [7], plants [8], or animals [9]. GWAS, particularly in humans, has yielded the “missing heritability” problem [10], though it is also present in other organisms (e.g., [11, 12]). Missing heritability is the gap between known and predicted heritability. In other words, GWAS has identified many genetic loci for a wide range of traits, but these markers do not account for all of the observed traits, implying the existence of the other undiscovered genetic factors [13]. An example of this missing heritability is a complex disease in human (e.g., Alzheimer’s disease [14]), and disease resistance in crops [12]. Devising new approaches and proper tools may help to uncover this missing heritability.

Previous work on GS and GWAS has focused primarily on statistical models, including Best Linear Unbiased Prediction (BLUP) and its variants [13, 15–18]. However, machine learning methods, such as random forests [19] and Support Vector Machines (SVMs) [20], have also seen an increasing interest in GS research on plants [21–26] and animals [27–29]. Also, random forests have been applied to GWAS on human or simulated data to identify markers that influence disease [30–33]. In this research, we employ sparse Bayesian learning [34] for predicting phenotypes and identifying influential markers on growth in the yeast *Saccharomyces cerevisiae*. This learning method uses Bayesian inference to obtain sparse solutions for regression and classification tasks. It is also called the Relevance Vector Machine (RVM), as it can be viewed as a kernel-based model of identical form to the SVM, which is a theoretically well-motivated classification algorithm in modern machine learning [35]. Although the prediction performance of RVMs practically competes with SVMs, they have some advantages that SVMs lack, such as having probabilistic outputs and the ability to work with arbitrary kernel functions. More importantly, RVMs construct much sparser models based on identifying more meaningful representatives of training data compared to the SVMs [34]. We use these representatives to help link phenotype predictions and identification of important markers in the yeast genome.

In this work, we consider the association problem as an embedded feature ranking problem wherein features are biological markers (e.g., SNPs), and the feature selection process is part of the predictive model construction. Then, the ranks of features based on their relevance to the trait will give candidate markers which can be further investigated in a GWAS. Motivated by the sparse solution property of sparse Bayesian learning, we investigate a novel ensemble architecture for feature selection and ranking. More precisely, we merge sparse Bayesian learning, ensemble and bagging techniques for ranking influential SNP markers on a quantitative trait. Note that there are also limited studies that used sparse Bayesian method for feature selection in bioinformatics [36–39]. However, this work, specifically on genes associated with disease, was only for classification, and did not incorporate ensemble techniques.

## Data and Methods

### Dataset

Bloom et al. [13] developed 1,008 haploid strains of *Saccharomyces cerevisiae* as a result of crosses between laboratory and wine strains of the yeast. The parent strains had sequence level differences of 0.5%. The genotypes consist of SNP markers that correspond to 11,623 sequence locations in the genome. The locations are coded as 1 if the sequence variation came from the wine strain parent, or 0 if it came from the laboratory strain parent.

Bloom et al. modified the environment of 1,008 yeast strains in 46 different ways (first column in Table 1), and measured the population growth under those different conditions. For example, they varied the basic chemicals used for growth (e.g. galactose, maltose), or added minerals (e.g. copper, magnesium chloride), then they measured growth in that condition. Precisely, Bloom et al. calculated the radius of the colonies from an image taken after approximately 48 hours of growth. Some results, such as irregular colonies, were removed and treated as missing data. Most traits have more than 90% of readings included.

**Table 1.** Coefficient of determination (*R*^2^) and standard deviation (std) of RVM predictions among the 46 traits.

| Trait | Linear Basis | Linear | Gaussian | Best RVM | std |
| --- | --- | --- | --- | --- | --- |
| Cadmium Chloride | 0.639 | 0.033 | 0.454 | Linear Basis | 0.005 |
| Caffeine | 0.074 | 0.216 | 0.233 | Gaussian($\gamma_3$) | 0.006 |
| Calcium Chloride | 0.113 | 0.273 | 0.287 | Gaussian($\gamma_3$) | 0.007 |
| Cisplatin | 0.133 | 0.29 | 0.287 | Linear | 0.006 |
| Cobalt Chloride | 0.258 | 0.439 | 0.466 | Gaussian($\gamma_2$) | 0.006 |
| Congo red | 0.327 | 0.467 | 0.491 | Gaussian($\gamma_1$) | 0.006 |
| Copper | 0.146 | 0.334 | 0.379 | Gaussian($\gamma_3$) | 0.01 |
| Cycloheximide | 0.317 | 0.473 | 0.514 | Gaussian($\gamma_1$) | 0.005 |
| Diamide | 0.277 | 0.473 | 0.483 | Gaussian($\gamma_2$) | 0.005 |
| E6 Berbamine | 0.211 | 0.375 | 0.414 | Gaussian($\gamma_2$) | 0.008 |
| Ethanol | 0.276 | 0.457 | 0.476 | Gaussian($\gamma_2$) | 0.006 |
| Formamide | 0.114 | 0.207 | 0.25 | Gaussian($\gamma_2$) | 0.006 |
| Galactose | 0.076 | 0.206 | 0.241 | Gaussian($\gamma_3$) | 0.008 |
| Hydrogen Peroxide | 0.234 | 0.343 | 0.397 | Gaussian($\gamma_2$) | 0.01 |
| Hydroquinone | 0.087 | 0.139 | 0.208 | Gaussian($\gamma_3$) | 0.009 |
| Hydroxyurea | 0.12 | 0.296 | 0.342 | Gaussian($\gamma_2$) | 0.01 |
| Indoleacetic Acid | 0.128 | 0.255 | 0.313 | Gaussian($\gamma_2$) | 0.007 |
| Lactate | 0.36 | 0.542 | 0.555 | Gaussian($\gamma_2$) | 0.005 |
| Lactose | 0.374 | 0.553 | 0.574 | Gaussian($\gamma_2$) | 0.008 |
| Lithium Chloride | 0.531 | 0.597 | 0.678 | Gaussian($\gamma_1$) | 0.006 |
| Magnesium Chloride | 0.102 | 0.245 | 0.255 | Gaussian($\gamma_3$) | 0.005 |
| Magnesium Sulfate | 0.187 | 0.366 | 0.41 | Gaussian($\gamma_3$) | 0.005 |
| Maltose | 0.409 | 0.484 | 0.523 | Gaussian($\gamma_2$) | 0.005 |
| Mannose | 0.079 | 0.213 | 0.197 | Linear | 0.007 |
| Menadione | 0.216 | 0.389 | 0.411 | Gaussian($\gamma_3$) | 0.006 |
| Neomycin | 0.422 | 0.583 | 0.596 | Gaussian($\gamma_2$) | 0.003 |
| Paraquat | 0.31 | 0.442 | 0.454 | Gaussian($\gamma_2$) | 0.005 |
| Raffinose | 0.185 | 0.385 | 0.388 | Gaussian($\gamma_3$) | 0.007 |
| SDS | 0.199 | 0.36 | 0.398 | Gaussian($\gamma_2$) | 0.004 |
| Sorbitol | 0.176 | 0.343 | 0.364 | Gaussian($\gamma_3$) | 0.009 |
| Trehalose | 0.326 | 0.48 | 0.503 | Gaussian($\gamma_2$) | 0.005 |
| Tunicamycin | 0.417 | 0.594 | 0.622 | Gaussian($\gamma_1$) | 0.006 |
| 4-Hydroxybenzaldehyde | 0.23 | 0.34 | 0.367 | Gaussian($\gamma_2$) | 0.008 |
| 4NQO | 0.44 | 0.496 | 0.512 | Gaussian($\gamma_2$) | 0.005 |
| 5-Fluorocytosine | 0.215 | 0.323 | 0.378 | Gaussian($\gamma_2$) | 0.008 |
| 5-Fluorouracil | 0.326 | 0.505 | 0.559 | Gaussian($\gamma_2$) | 0.005 |
| 6-Azauracil | 0.152 | 0.3 | 0.304 | Gaussian($\gamma_3$) | 0.005 |
| Xylose | 0.282 | 0.455 | 0.478 | Gaussian($\gamma_3$) | 0.004 |
| YNB | 0.379 | 0.224 | 0.515 | Gaussian($\gamma_1$) | 0.009 |
| YNB:ph3 | 0.059 | 0.18 | 0.177 | Gaussian($\gamma_3$) | 0.005 |
| YNB:ph8 | 0.203 | 0.327 | 0.361 | Gaussian($\gamma_2$) | 0.006 |
| YPD | 0.368 | 0.266 | 0.511 | Gaussian($\gamma_1$) | 0.008 |
| YPD:15C | 0.211 | 0.334 | 0.356 | Gaussian($\gamma_2$) | 0.006 |
| YPD:37C | 0.473 | 0.566 | 0.611 | Gaussian($\gamma_2$) | 0.006 |
| YPD:4C | 0.18 | 0.406 | 0.438 | Gaussian($\gamma_2$) | 0.005 |
| Zeocin | 0.316 | 0.46 | 0.475 | Gaussian($\gamma_3$) | 0.004 |

Gaussian parameter: $\gamma_1 = 1e-4$ , $\gamma_2 = 2e-4$ , and $\gamma_3 = 3e-4$ .

### Sparse Bayesian Learning

The sparse Bayesian modelling [34, 40] is an approach for learning the prediction function *y*(**x; w**), which is expressed as a linear combination of basis functions:

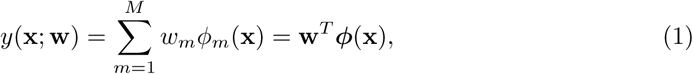
where *ϕ*(**x**) = (*ϕ*(**x**),…,*ϕ_M_*(**x**))*^T^* are basis functions, generally non-linear, and *w*_1_, *…,w_M_* are the adjustable parameters, called weights. Given a dataset of input-target training pairs 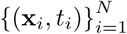, the objective of the sparse Bayesian method is to estimate the target function *y*(**x; w**), while retaining as few basis functions as possible. The sparse Bayesian algorithm often generates exceedingly sparse solutions (i.e., few non-zero parameters *w_i_*).

In a particular specialization of (1), such as the one that SVM uses, *M* = *N* and the basis functions take the form of kernel functions, one for each data point **x***_m_* in the training set, so that *ϕ_m_*(**x**) = *K(***x**, **x***_m_*), where *K*(.,.) is the kernel function. This exemplification of the sparse Bayesian modelling is called the Relevance Vector Machine (RVM). Tipping [41] introduced the RVM method as an alternative to the SVM method of Vapnik [42]. However unlike SVMs, where the kernel functions must be Positive Definite Symmetric (PDS) [43], we can use arbitrary basis sets in the RVM.

Assuming that the basis functions have the form of kernel functions, we illustrate the sparse Bayesian algorithm for regression in the following. Corresponding algorithms for arbitrary basis functions can be easily induced from them.

#### Relevance Vector Regression

We follow the framework developed by Tipping [34]. In the regression framework, the targets **t** = (*t*_1_,…,*t_N_*)*^T^* are real-valued labels. Each target *t_i_* is representative of the true model *y_i_*, but with the addition of noise *ε_i_*: *t_i_* = *y*(*x_i_*) + *ε_i_*, where *ε_i_* ~ *N* (0, *σ*^2^). This means *p(t_i_* | **x***_i_*, **w**, *σ*^2^) = *N(y(***x***_j_), σ^2^)*, or

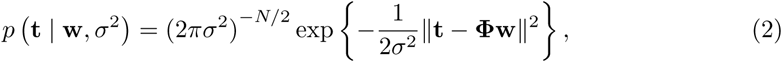
where **w** = (*w*_1_,…, *w_N_*)*^T^*, and the data is hidden in the design matrix (kernel matrix) **Φ** = [*ϕ*(**x**_1_),…, *ϕ*(*x_N_*)], wherein *ϕ*(**x***_i_*) = [*K*(**x***_i_*, **x**_1_),…, *K*(**x***_i_*, *x_N_*)]. For clarity, we omit the implicit conditioning on the set of input vectors {**x***_i_*} in (2) and subsequent expressions.

We infer weights using a fully probabilistic framework. Specifically, we define a Gaussian prior distribution with zero mean and *α_i_*^−1^ variance over each ***w****_i_*: *p* (*w_i_* | *α_i_*) = *N* (0, *α_i_*^−1^), or:

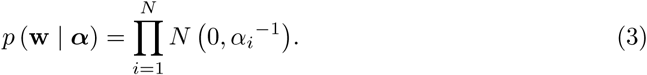

The sparsity of the RVM is a result of the the independence of the hyperparameters ***α*** = (*α*_1_,…, **α***_N_*)*^T^* one per basis function (i.e., weight), which moderate the strength of the prior information [44]. Using Bayes’ rule and having the prior distribution and likelihood function (3) and (2), the posterior distribution over the weights would be a multivariate Gaussian distribution:

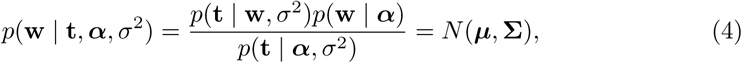
where the covariance and the mean are:

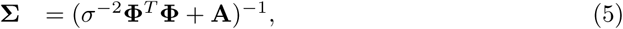

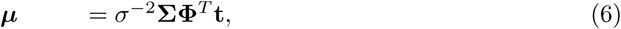
and **A** = *diag (α_1_,…, α_N_*).

The likelihood distribution over the training target **t**, given by (2), is marginalized with respect to the weights to obtain the marginal likelihood for the hyperparameters:

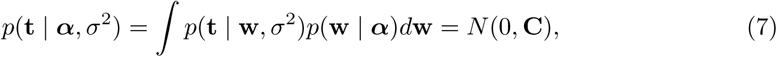
where the covariance is given by 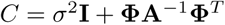. Values of ***α*** and *σ^2^* which maximize (7) cannot be obtained in closed form, thus the solution is derived via an iterative maximization of the marginal likelihood *p*(**t** | ***α****, σ*^2^) with respect to ***α*** and σ^2^:

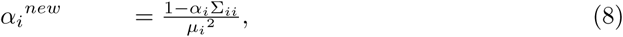

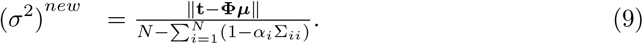

By iterating over (5), (6), (8), and (9), the RVM algorithm reduces the dimensionality of the problem when *α_i_* is larger than a threshold (note that *α_i_* has a negative power in (3)) [45]. The algorithm stops when the likelihood *p*(**t** | ***α****, σ^2^)* stops increasing. The non-zero elements of **w** are called Relevance Values. The input vectors which correspond to the relevance values are called Relevance Vectors (RVs) as an analogy to Support Vectors in the SVM [45]. Having the relevance vectors, 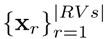, and the relevance values, 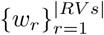, the RVM makes prediction on a new data instance **x**_*_:

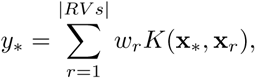

where |*RVs*| denotes the cardinality of the set of relevance vectors.

The regression framework can be extended to the classification case using the approximation procedure presented in [41].

### Kernel RVM versus Basis RVM

Kernel methods are flexible techniques that can be used to extend algorithms such as SVMs to define non-linear decision boundaries [46]. For example, consider a binary classification problem in which input patterns are not linearly separable in the input space (i.e., inputs cannot be separated into two classes with passing a hyperplane between them). In such a case, one solution is to use a non-linear mapping of the inputs into some higher-dimensional feature space in which the patterns are linearly separable. Then, we solve the problem (i.e., finding the optimal hyperplane) in the feature space, and consequently, we will be able to identify the corresponding non-linear decision boundary for the input vectors in the input space. To do this procedure, a kernel method only requires a function *K*: *X* × *X* → R, which is called a kernel over the input space *X*. For any two input patterns **x***_i_*, **x***_j_* ∈ *X*, *K*(**x***_i_*, **x***_j_*) is the dot product of vectors *ϕ*(**x***_i_*) and *ϕ*(**x***_j_*) for some mapping *ϕ*: *X* → *H* to a feature space *H*:

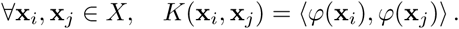

In this research, we define sparse Bayesian learning in such a way that we can discriminate between kernel and basis functions, i.e., “kernel” RVM versus “basis” RVM. The basis RVMs, which do not have counterparts in SVMs, will be mainly used to enable feature selection. For example, we define two types of linear RVMs, which we call linear kernel RVMs and linear basis RVMs. In a linear kernel RVM, the basis functions in (1) are linear kernel functions, i.e.,

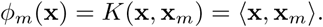

When we use linear kernels, in fact we have no mapping. In other word, there is no feature space (as we use input vectors directly), so our estimator tries to pass a hyperplane through input vectors in the input space (e.g., in the case of regression).

In our linear basis RVM, the basis functions are linear and equal to the features of the input vectors, i.e.,

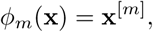

where **x**^[^*^m^*^]^ refers to the *m*-th feature in an input vector **x** with *M* dimensions. We can view it as if we have no basis function in a linear basis RVM, as we use input vectors directly in (1) instead:

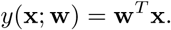

Therefore, we can restate (2) with weights **w** =(*w*_0_*,w*_1_*,…, w_M_*)*^T^*, where *M* is the number of features, and the design matrix is

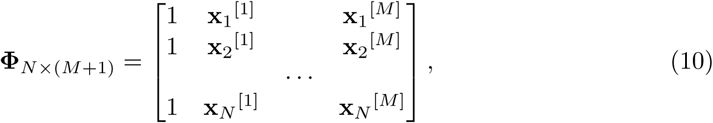
where the first column handles the intercept *w*_0_, and *N* is the number of training individuals.

Thus, this linear basis RVM will find the RVs which corresponds to the features; i.e., the obtained sparsity will be in the feature set rather than the training individuals. This is exactly what we expect from a feature selection method. Therefore, this RVM can perform target prediction as well as feature selection. For example, in a GS in crop breeding, the individuals are breeds of a crop, the features are the markers (SNPs), and a phenotype is a target. Then, a linear basis RVM would identify a subset of relevant markers to that phenotype, while it is trained for phenotype prediction.

Similar to linear RVMs, we can define any other non-linear RVMs (i.e., Gaussian RVM as Gaussian kernel RVM or Gaussian basis RVM). In our experiments, we apply kernel RVMs with different PDS kernel types to investigate how they perform in predicting phenotypes. However, we only examine linear basis RVMs for phenotype prediction and influential marker identification.

Compared to the SVM method, we should note that there is not an SVM counterpart for a basis RVM, as the design matrix (10) resembles a non-PDS function which specifically cannot be used in an SVM. In a kernel RVM, we can use PDS kernels, such as polynomial and Gaussian kernels, or non-PDS kernels, such as sigmoid kernels (neural network kernels [47]). In the case of using PDS kernels, the kernel RVM prediction accuracies will be comparable to the SVM results.

#### Kernel Types

In our experiments with kernel RVMs, we use both sequence and non-sequence kernel functions. A non-sequence kernel refers to a kernel that can handle binary or numerical data types (e.g., gene expression data). Gaussian kernel and polynomial kernel are among non-sequence kernels: For any constant *γ >* 0, Gaussian kernel is the kernel *K*: ℝ*_N_* → ℝ:

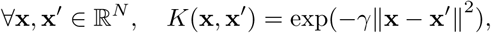

where 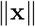 is the norm of the vector **x**. Also, a polynomial kernel of degree *d* such as *K* is defined by:

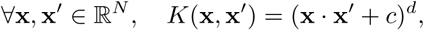

for a fixed constant *c* ≥ 0. A linear kernel is a polynomial kernel with *c* = 0 and *d* = 1.

In contrast to a non-sequence kernel, a sequence kernel operates on strings, or finite sequences of symbols. Intuitively speaking, we can say that the more similar the two strings **x** and **y** are, the higher the value of a string kernel *K*(**x**, **y**) will be. The *n*-gram kernel [48] is an example of a sequence kernel. The *n*-gram kernel of the two strings **x** and **y** counts how often each contiguous string of length *n* is contained in the strings:

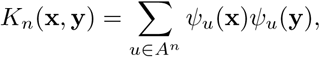

where *ψ_u_*(**x**) denotes the number of occurrences of the subsequence *u* in the string **x**, and *A_n_* is the set of all possible subsequence of length *n*, given the alphabet *A*. For instance, suppose we are given two DNA sequences with the alphabet *A* = {A,C,G,T}: **x** = AACCT and **y** = GACAC. The bi-gram (2-gram) subsequences in **x** and **y** are {AA,AC,CC,CT} and {GA,AC,CA}, respectively. Therefore, *K*_2_(**x**, **y**)=1 × 2=2, as only one subsequence AC is common in both sequences, which it has been repeated once in **x** and twice in **y**. Higher kernel values mean two sequences are more similar.

The sequence kernels used in applications such as computational biology are rational kernels [49]. Rational kernels [46, 50], which are based on finite-state transducers [51], present an efficient general algorithm for manipulating variable-length sequence data. For computing rational kernels, we use OpenFST (Open Finite-State Transducer) library [52, 53] and OpenKernel library [54].

#### RVM as a Phenotype Predictor

We consider the yeast dataset as 46 separate regression problems: we construct a separate RVM model for predicting growth under each of 46 conditions. We train each RVM with linear basis function, linear kernel, Gaussian kernel (with different values of *γ* parameter), and a set of *n*-gram kernels. Using the coefficient of determination (*R*^2^) as measure, and running 10 times of 10-fold cross-validation (each time with random different folds), we evaluate the results of RVM models. As the process for this dataset along with repeating cross-validations is computationally heavy, the process is done in parallel on the WestGrid (www.westgrid.ca) platform.

### Ensemble RVM

In an ensemble, a set of classifiers is trained and for new predictions, the results of each of the classifiers is combined to obtain a final result [55]. Ensembles are often produce better predictive performance than a single model by decreasing variance (bagging), bias (boosting), or improving predictions (stacking) [56]. Moreover, ensemble techniques have the advantage of handling large data sets and high dimensionality because of their divide-and-conquer strategy. Random Forests [19] and Gradient Boosting Machines (GBMs) [57] are examples of ensemble methods.

In this research, we employ ensemble RVM with bagging approach. Bagging (bootstrap aggregating [58]) is based on bootstrapping, where sample subsets of a fixed size are drawn with replacement from an initial set of samples. In bagging, a large number of separate classifiers in an ensemble are trained on separate bootstrap samples and their predictions are aggregated through majority voting or averaging. Bagging is commonly used as a resolution for the instability problem in estimators.

We use ensembles of basis RVMs for feature selection and ranking. Each RVM model in an ensemble finds a set of representatives (the RVs) which represent important features. Then, aggregating RVs of the ensemble lets rank the features. The top ranked markers are chosen based on a threshold. In other words, we define the most influential markers as those who are chosen by a specific percentage of the RVMs in the ensemble as RVs. Ranking mechanisms allow us to reduce dimensionality and enhance generalization [59]. Furthermore, they enable us to recognize interpretable or insightful features in the model.

We use SpareBayes software package for Matlab [60] to implement the RVMs in this research.

## Results and Discussion

### Predicting Phenotypes

The prediction accuracies plus the standard deviation of cross-validation results in the best RVM model are shown in Table 1. The value reported for the *γ* parameter of Gaussian function in the table is the best of a range of values we tried for model selection. Note that Gaussian kernel RVMs mostly produce promising results. Even in traits such as Cisplatin and Mannose, the linear kernel RVM shows a slightly better accuracy than the Gaussian. The only exception is Cadmium Chloride in which linear basis RVM presents a significantly better accuracy. The RVM models are stable, based on the standard deviations. In following subsections, we analyse the results with more details.

#### Linear Kernel RVM versus Linear Basis RVM

As explained before, a linear basis RVM can be viewed as an RVM with no basis function, as we use input vectors directly in the data model instead. Similarly when we use linear kernels, it means we do not map the inputs into a higher dimensional feature space, so our estimator tries to pass a hyperplane through input vectors in the input space. Here, we might expect that both linear kernel and linear basis RVMs produce similar results or with subtle difference, as both are linear and in the same space. However, that is not the case, i.e., linear kernel RVM and linear basis RVM produces different hyperplanes as we see in the results in Table 1. Consider Cadmium Chloride and YPD:4C, as two extreme examples. In the former, the linear basis RVM has high accuracy, while in the latter the linear kernel RVM shows higher accuracy. As a corollary we can say that linear basis RVM produces results which classic linear SVM is not able to. We know that the linear kernel cannot be more accurate than a properly tuned Gaussian kernel [61], but we cannot conclude the same for the linear basis function. Therefore, even if we have conducted a complete model selection using the Gaussian kernel RVM for a problem, it is still valuable to consider the linear basis RVM, just as we saw linear basis superiority to Gaussian kernel in Cadmium Chloride.

#### Investigating String Kernel RVM

We have also investigated the *n*-gram kernel, a form of string kernel, with the RVM, for *n* =3, 5, 7, 10. All string kernels showed poor accuracies on our dataset. The issue arises from the fact that a typical *n*-gram kernel on this dataset gives us a kernel matrix with almost all elements close to one. It intuitively indicates that the sequences are so similar to each other that the predictor cannot discriminate between any pairs. One possible explanation for the poor performance of the *n*-gram kernels is genetic linkage. Genetic linkage describes an inheritance tendency in which two markers located in close proximity to each other on the same chromosome are more likely to be inherited together during meiosis [62]; i.e, the nearer two genes are on a chromosome, the lower the chance of recombination between them, and the more likely they are to be inherited together. *N*-gram kernels capture the short adjacent similarities in sequences. Therefore, high similarity between sequences captured by *n*-gram kernels comes as no surprise. That is, we expect the small 3-10 SNP sequences to be shared between individuals because these sequences appear close to each other in the genome and are similar due to genetic linkage. The genetic linkage phenomenon can also illustrate why *n*-gram kernels previously helped for gene-scale problems such as metabolic network prediction [63], but do not work for this problem which has a genome-scale attribute.

#### Heritability versus Accuracies

Bloom et al. [13] provided estimates for narrow-sense and broad-sense heritability for the yeast dataset. They considered broad-sense heritability as the contribution of additive genetic factors (i.e., narrow-sense heritability) and gene-gene interactions. Thus, the broad-sense heritability is always greater than the narrow sense heritability, and their difference can be interpreted as a measurement of gene-gene interactions [13]. The broad-sense heritability estimates among the 46 traits ranged from 0.40 (YNB:ph3) to 0.96 (Cadmium Chloride), with a median of 0.77. Also, the narrow-sense heritability estimates ranged from 0.21 (YNB:ph3) to 0.84 (Cadmium Chloride), with a median of 0.52. Using the difference between two heritability measures, Bloom et al. estimated the fraction of genetic variance due to gene-gene interactions, which ranged from 0.02 (5-Fluorouracil) to 0.54 (Magnesium Sulfate), with a median of 0.30. Therefore, the genetic basis for variation in some traits, such as 5-Fluorouracil, is almost entirely due to additive effects, while for some others, such as Magnesium Sulfate, approximately half of the heritable component is due to gene-gene interactions.

To determine if there is a correlation between heritability and RVM prediction accuracies, we calculated the Pearson correlation coefficient between estimates of heritability and prediction accuracies. The correlation coefficients in three RVM categories (Gaussian, linear, and linear basis) are shown in Fig 1. The values related to the broad-and narrow-sense heritability (blue and orange bars) indicate that heritability and RVM accuracies, particularly in Gaussian and linear basis RVMs, have strong positive association. In other words, we will have better predictions when the amount of heritability increases. In particular, a higher narrow-sense heritability yields better prediction rates for the RVM predictor.

**Fig 1.**
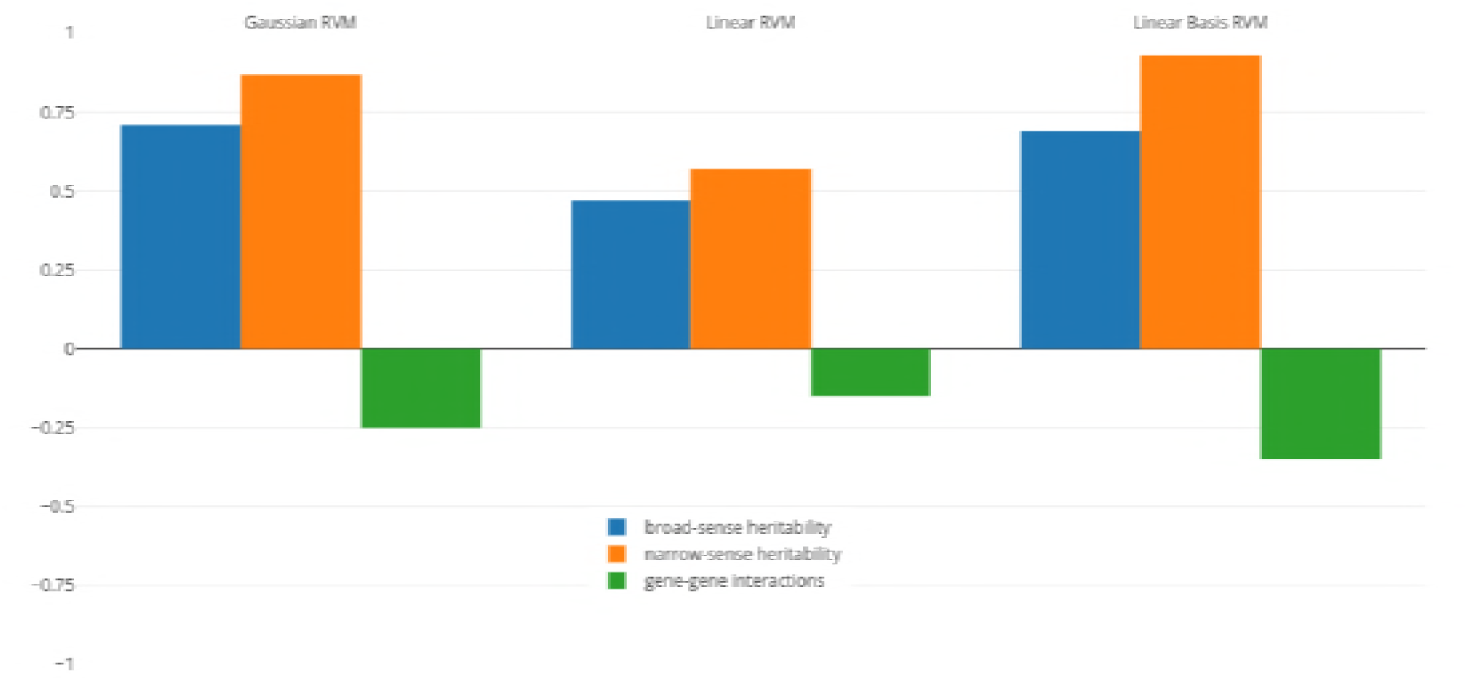
Pearson correlation coefficient between RVM accuracies and different heritability measures.

To determine if RVMs are less successful in predicting traits with larger non-additive effects, we also calculated the correlation coefficient between RVM accuracies and gene-gene interactions effects (green bars in the figure). These values indicate that gene-gene effects and accuracies, particularly in Gaussian and linear RVMs, have small negative association, indicating that we cannot infer the RVM performance is deteriorating when gene-gene interactions effects increases. This confirms previous results where non-parametric and semi-parametric machine learning techniques, such as SVMs, RKHS, and random forests, have been shown to have good prediction abilities for non-additive traits [64, 65]. However, if we have narrow-sense heritability estimates before constructing an RVM model, we are able to anticipate behaviour of the predictor, due to the higher weight of additive effects (as most genetic variance in populations is additive [66]).

#### Comparison with Related Work

Grinberg et al. [25] recently compared several learning methods including forward stepwise regression, ridge regression, lasso regression, random forest, GBM, and Gaussian kernel SVM with two classical statistical genetics methods (BLUP and a linkage analysis done by Bloom et al. [13]). Grinberg et al. used the coefficient of determination (*R*^2^) as accuracy measure, and evaluated their models with one run of 10-fold cross validation. In Table 2, the columns “G: Best of Others” and “G: SVM” refer to Grinberg et al.’s results. Also, the *R*^2^ value in the RVM column belongs to the best RVM given in Table 1.

**Table 2.** RVM results versus Grinberg et al.’s (G) [25].

| Trait | G: Best of Others | G: SVM | RVM(std) |
| --- | --- | --- | --- |
| Cadmium Chloride | GBM:0.797 | 0.565 | 0.639(0.005) |
| Caffeine | GBM:0.250 | 0.234 | 0.233(0.006) |
| Calcium Chloride | BLUP: 0.268 | 0.261 | 0.287(0.007) |
| Cisplatin | GBM: 0.338 | 0.272 | 0.287(0.006) |
| Cobalt Chloride | GBM: 0.460 | 0.448 | 0.466(0.006) |
| Congo red | Lasso:0.504 | 0.487 | 0.491(0.006) |
| Copper | GBM:0.452 | 0.338 | 0.379(0.01) |
| Cycloheximide | SVM:0.529 | 0.529 | 0.514(0.005) |
| Diamide | BLUP:0.498 | 0.486 | 0.483(0.005) |
| E6 Berbamine | GBM:0.412 | 0.390 | 0.414(0.008) |
| Ethanol | GBM:0.518 | 0.455 | 0.476(0.006) |
| Formamide | GBM:0.350 | 0.240 | 0.25(0.006) |
| Galactose | GBM:0.235 | 0.217 | 0.241(0.008) |
| Hydrogen Peroxide | SVM:0.399 | 0.399 | 0.397(0.01) |
| Hydroquinone | BLUP:0.225 | 0.188 | 0.208(0.009) |
| Hydroxyurea | GBM:0.337 | 0.301 | 0.342(0.01) |
| Indoleacetic Acid | Bloom et al.:0.480 | 0.3 | 0.313(0.007) |
| Lactate | Lasso:0.568 | 0.557 | 0.555(0.005) |
| Lactose | GBM:0.582 | 0.565 | 0.574(0.008) |
| Lithium Chloride | GBM:0.711 | 0.680 | 0.678(0.006) |
| Magnesium Chloride | Bloom et al.:0.278 | 0.267 | 0.255(0.005) |
| Magnesium Sulfate | Bloom et al.:0.519 | 0.378 | 0.41(0.005) |
| Maltose | GBM:0.809 | 0.522 | 0.523(0.005) |
| Mannose | GBM:0.255 | 0.215 | 0.213(0.007) |
| Menadione | GBM:0.432 | 0.402 | 0.411(0.006) |
| Neomycin | Lasso:0.614 | 0.597 | 0.596(0.003) |
| Paraquat | Lasso:0.496 | 0.479 | 0.454(0.005) |
| Raffinose | GBM:0.383 | 0.364 | 0.388(0.007) |
| SDS | Lasso:0.411 | 0.383 | 0.398(0.004) |
| Sorbitol | Bloom et al.:0.424 | 0.318 | 0.364(0.009) |
| Trehalose | GBM:0.515 | 0.477 | 0.503(0.005) |
| Tunicamycin | SVM:0.634 | 0.634 | 0.622(0.006) |
| 4-Hydroxybenzaldehyde | GBM:0.397 | 0.36 | 0.367(0.008) |
| 4NQO | GBM:0.636 | 0.542 | 0.512(0.005) |
| 5-Fluorocytosine | GBM:0.399 | 0.364 | 0.378(0.008) |
| 5-Fluorouracil | Lasso:0.552 | 0.546 | 0.559(0.005) |
| 6-Azauracil | GBM:0.315 | 0.279 | 0.304(0.005) |
| Xylose | GBM:0.516 | 0.460 | 0.477(0.004) |
| YNB | GBM:0.543 | 0.525 | 0.515(0.009) |
| YNB:ph3 | BLUP:0.195 | 0.166 | 0.177(0.005) |
| YNB:ph8 | BLUP:0.356 | 0.334 | 0.361(0.006) |
| YPD | GBM:0.556 | 0.524 | 0.511(0.008) |
| YPD:15C | Bloom et al.:0.432 | 0.333 | 0.356(0.006) |
| YPD:37C | Bloom et al.:0.711 | 0.603 | 0.611(0.006) |
| YPD:4C | GBM:0.485 | 0.421 | 0.438(0.005) |
| Zeocin | GBM:0.495 | 0.475 | 0.472(0.004) |

Compared to the SVM, RVM models show better predictions overall. However, Grinberg et al.’s approach for training and model selection in Gaussian SVM is not proper. The authors trained an SVM with distinct parameters for each fold of cross-validation. In other words, they trained 10 SVMs (10 sets of Gaussian kernel and SVM parameters) for a trait. In this way, not only the accuracies are overestimated, but also the model selection process appears problematic (e.g., the set of parameters that should be used to predict a trait for new yeast individuals is unclear).

The RVM is comparable to the best of the methods tested by Grinberg et al., except in six traits including Cadmium Chloride, Indoleacetic Acid, Magnesium Sulfate, Maltose, 4NQO, and YPD:37C in which GBM or Bloom et al.’s method showed superiority. However, the mean broad sense heritability of these six traits is 0.88, and the mean narrow sense heritability is 0.66. This confirms that nonlinear techniques, including GBM and RVM, are competitive for predictions involving traits with high broad sense heritability. Also, we should note that we do not know about the stability of the methods experimented by Grinberg et al., as they ran only one 10-fold cross-validation, while the RVM shows high stability, as its standard deviations in 10 runs of 10-fold cross-validation were small.

### Identifying Influential Markers

For identifying the most influential markers (SNPs) on the traits, we used our RVM ensemble architecture for ranking markers. An ensemble for a trait was composed of 400 linear basis RVMs, each with subsampling 50 to 60% of training data. As we are only interested in a small set of top ranked markers, we observed that the size of subsampling does not affect the results (data not shown). To demonstrate how well the ensemble RVMs act in identifying influential markers, we present the top ranked markers in three conditions (traits): Cadmium Chloride, Lithium Chloride, and Mannose. We chose Cadmium Chloride and Mannose as samples which the linear basis RVM showed excellent and poor phenotypic prediction accuracies (Table 1), respectively, while we chose Lithium Chloride for comparison to the work of Bloom et al. [13]. Also, these conditions are across a wide range of broad sense heritability: the broad sense heritability of Cadmium Chloride is 0.98, Mannose is 0.42 and Lithium Chloride is 0.87.

The ensemble RVMs for each of the three traits ranked around 90% of the markers with rank values in the range [1, 400]. The unranked markers indicate the markers that do not have any effect (even minor) on a trait. We define the most influential markers as those that are chosen by half of the RVMs in the ensemble as RVs, so in this dataset we will have less than ten influential markers in the three traits. The ranked markers indicate those who may have positive or negative effects on a trait. In other words, we not only find the markers which have additive effects on yeast growth in an environment, but also we find those which have adverse effects on growth.

#### Comparison with Related Work

Previously, Bloom et al. [13] conducted a linkage analysis with high statistical power to map functional QTL in all 46 traits. They found that nearly the entire additive genetic contribution to heritable variation (narrow-sense heritability) in yeast can be explained by the detected loci. Bloom et al. specifically showed that for one trait (Lithium Chloride), the loci detected by their method explained most of the heritability.

We compare our identified influential markers in three traits to Bloom et al.’s QTL. Bloom et al. found 6, 22, and 10 additive QTL in Cadmium Chloride, Lithium Chloride, and Mannose, respectively. Therefore, we chose the top 6, 22, 10 ranked SNPs in the three traits as well. Figs 2, 3 and 4 show results in each of the three traits accordingly. Each of the figures includes two parts (a) and (b) corresponding to the map of yeast chromosomes 1-8 and 9-16, respectively. The results were demonstrate that the markers identified by the RVM ensembles have similar distribution to the Bloom et al.’s QTL. Also, the RVM ensembles were relatively successful in finding the exact markers in the traits (33% match rate in Cadmium Chloride, 36% in Lithium Chloride, and 40% in Mannose). We note that the highest match rate among the three traits belongs to Mannose in which the linear basis RVM had poor prediction accuracy. This could be an advantage of the RVM being capable of recognizing true “representatives” of a population, despite unacceptable predictions. Another advantage is in the ranking system, where we can always recognize the effect of a marker on a trait with its weight, even in the small set of top-ranked markers. However, we can also go further and conclude that those top ranked markers that are close to each other (e.g, markers at loci 649 kb, 656 kb, and 677 kb on Chromosome 12 in Fig 3) suggest to a higher impact of a locus near to those markers on a trait due to genetic linkage.

**Fig 2.**
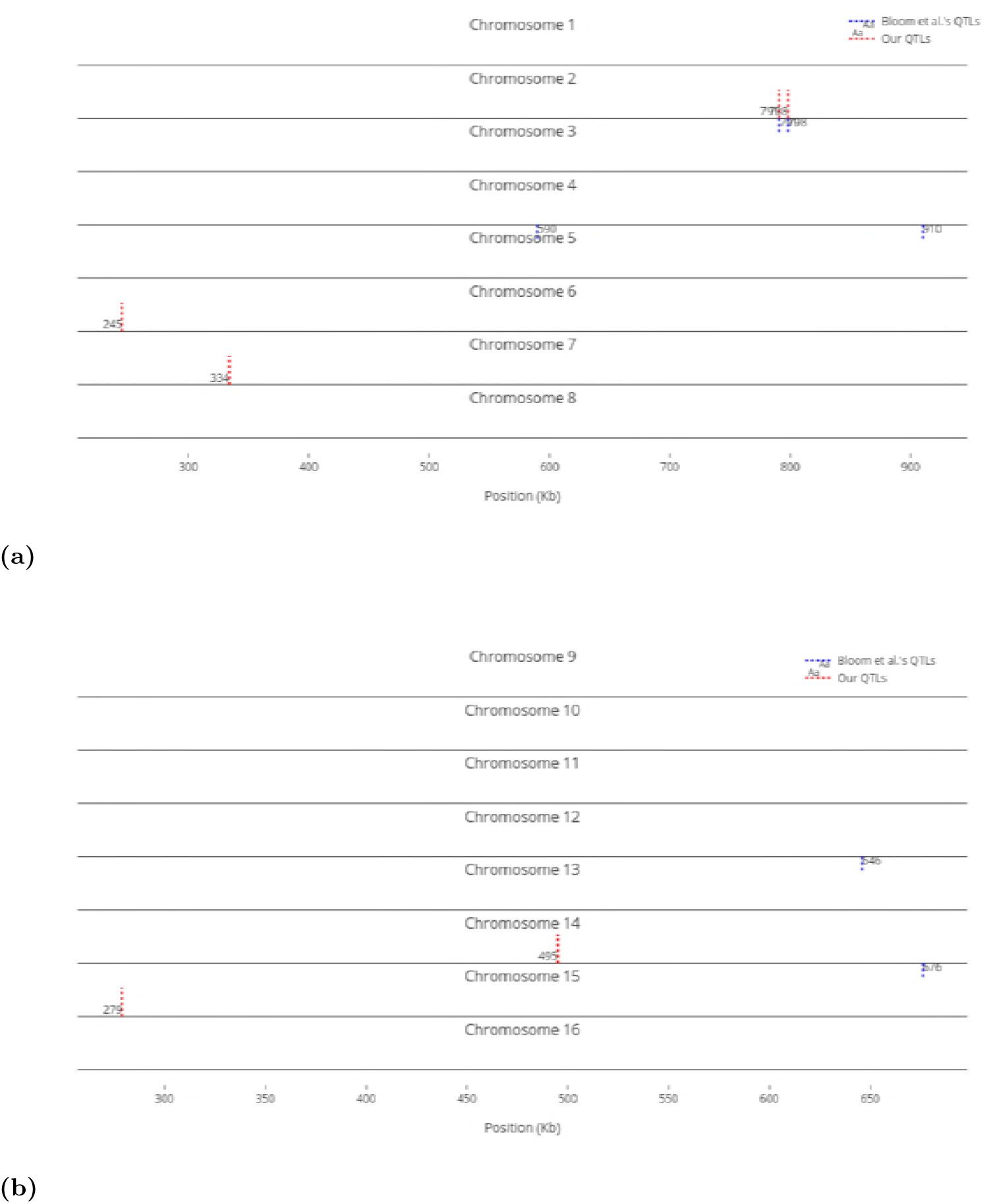
Top 6 influential markers on growth in Cadmium Chloride recognized by ensemble RVMs versus Bloom et al.’s 6 QTL.

**Fig 3.**
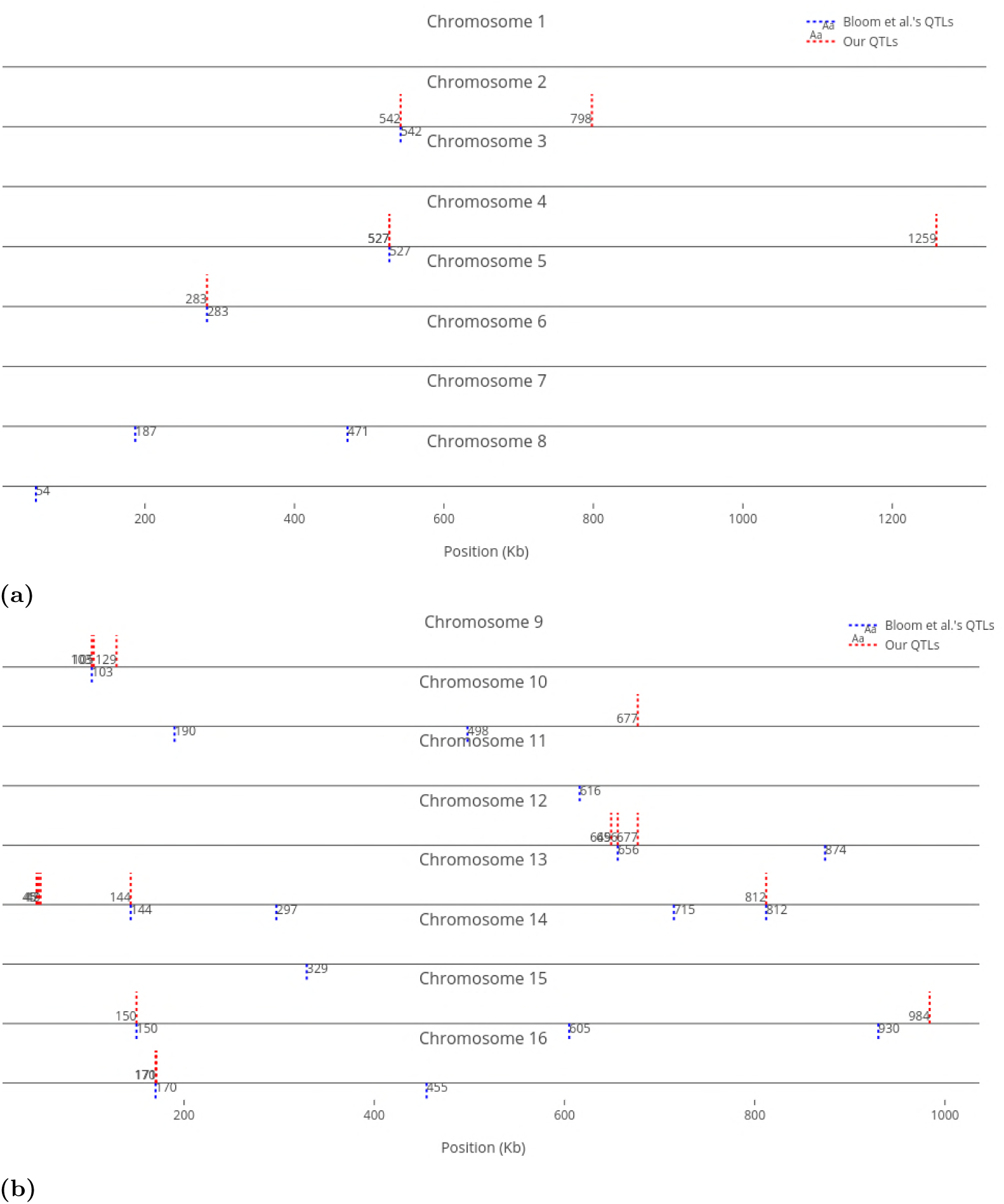
Top 22 influential markers on growth in Lithium Chloride recognized by ensemble RVMs versus Bloom et al.’s 22 QTL.

**Fig 4.**
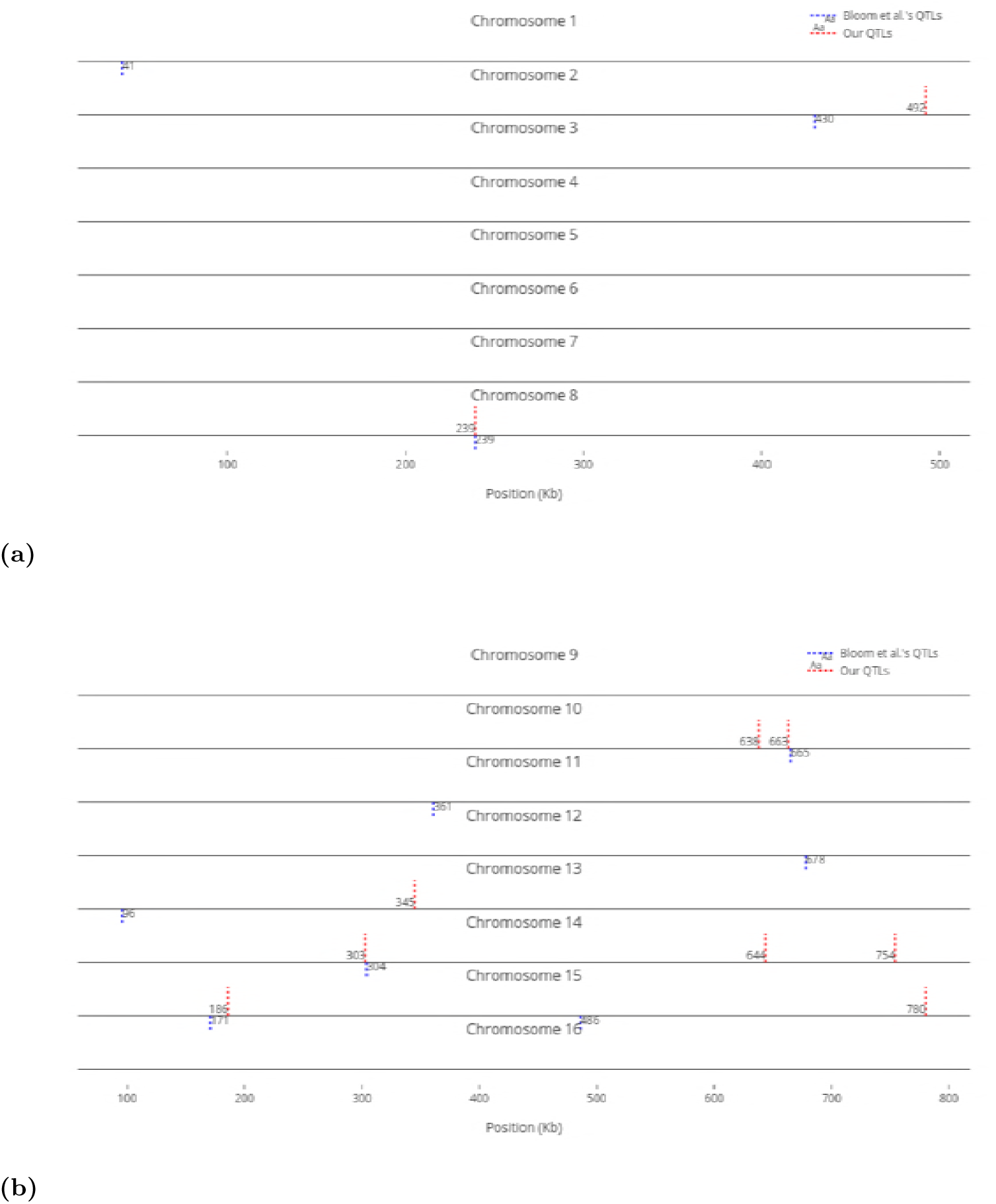
Top 10 influential markers on growth in Mannose recognized by ensemble RVMs versus Bloom et al.’s 10 QTL.

For comparison purposes, we only provided an equal number of top ranked markers to Bloom et al.’s QTL. However, when we decrease the threshold, the number of influential markers would increase. For instance, Fig 5 shows the top ten (instead of six) most influential markers in Cadmium Chloride. In this case, another additive QTL in chromosome 12 is identified (i.e., at position 464 kb). As not all influential markers have additive effects, the identified markers which are distant from Bloom et al.’s QTL present a good set of candidates for further investigation, to determine if they have non-additive effects with other loci.

**Fig 5.**
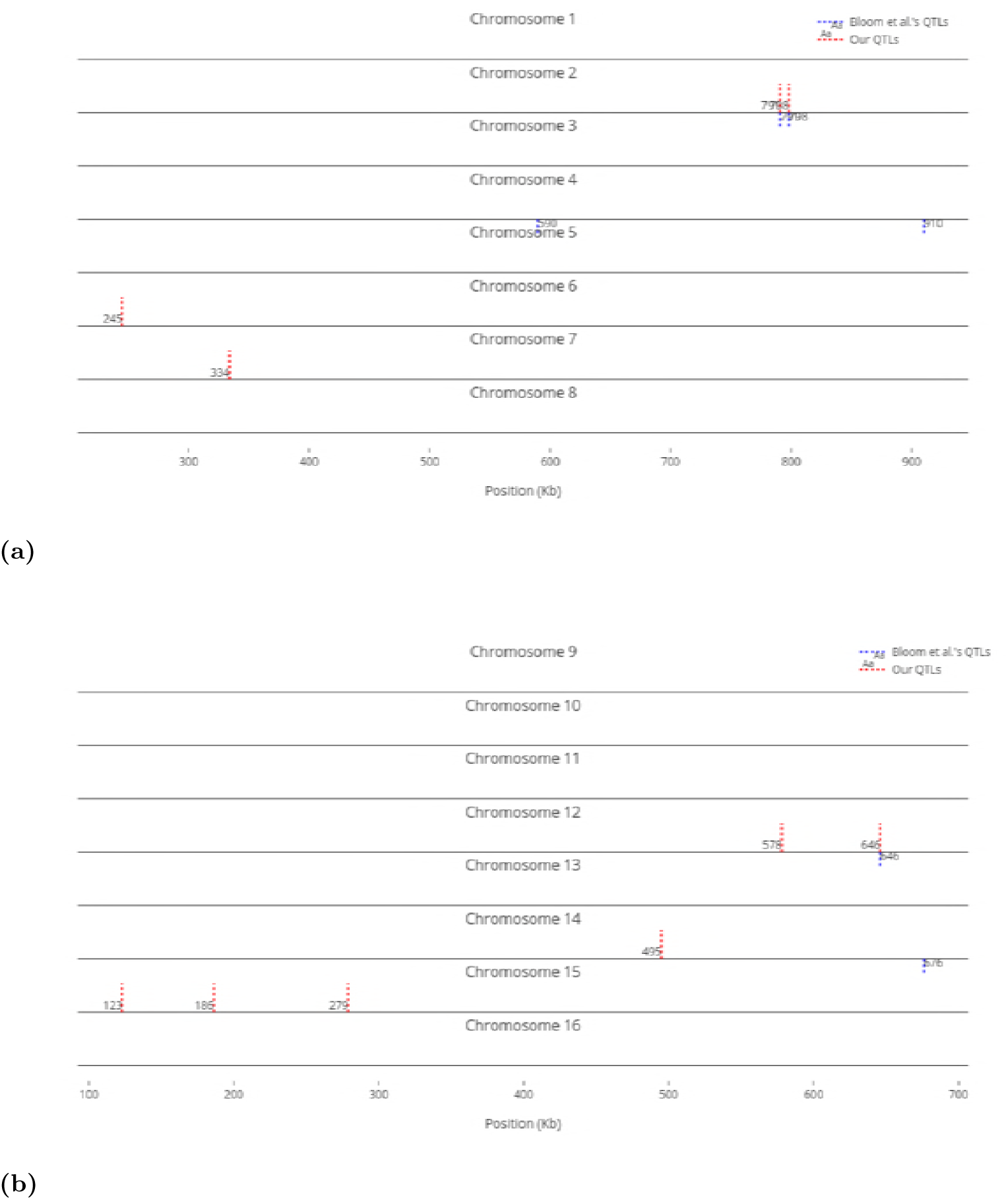
Top 10 influential markers on growth in Cadmium Chloride recognized by ensemble RVMs versus Bloom et al.’s 6 QTL.

## Conclusion

In this research, we studied how RVMs perform on growth prediction of yeast in 46 different environments, comparing its performance with other learning methods such as SVM and GBM. Our obtained phenotype prediction accuracies suggest that RVM shows positive results, and can be used as an alternative method in genomic selection. It is well-known that no machine learning technique performs best in all datasets [25, 67].

We investigated different kernels in RVM. We illustrated how different linear RVMs, i.e, linear kernel RVM and linear basis RVM, perform in phenotype prediction. We observed that Gaussian RVMs had the best accuracies, while string kernel RVM, such as *n*-gram, presented poor predictions.

We also investigated the relationship between different heritability measures and RVM prediction accuracies. The results indicate an strong association between narrow-sense heritability and prediction accuracy in RVMs. On the other hand, new research points out that the most genetic variance in populations is additive [66]. Therefore, if the heritability is known in advance, we can consequently anticipate the performance of the model before constructing it.

The last part of the experiments was devoted to identifying most influential markers on the traits, as well as non-relevant markers. We chose three traits with different phenotype prediction accuracies as samples, and demonstrated how well our RVM ensembles work to rank the markers in each trait, comparing the results with other research which used a traditional linkage analysis to find additive QTL. The comparison validates the results of RVM ensembles in finding markers with additive effects. However, we can earn more from the RVM ensembles, as those are capable of identifying both growth-increasing and growth-decreasing markers in yeast.

It may perhaps be observed that our ensemble linear basis RVM for feature selection takes in to account only linear relationships. Although this linear separability is a reasonable assumption for high dimensional data, it is desirable to investigate nonlinear basis substitution, particularly Gaussian functions, to handle nonlinear relationships. Gaussian basis RVM still gives feature RVs as each Gaussian basis in the model operates on a different dimension (feature). However, employing Gaussian basis RVM requires setting not only the variance (*σ_m_*) in each Gaussian basis function in (1), but also the mean or center (*µ_m_*): *ϕ_m_*(**x**) = exp(−(**x**^[^*^m^*^]^ − *µ_m_*)^2^*/σ_m_* ^2^), where **x**^[^*^m^*^]^ refers to the *m*-th feature in an input vector **x** with *M* dimensions. Investigating any appropriate approach, with acceptable computational complexity, for choosing parameters in Gaussian basis RVMs, and employing these RVMs in an application remain as future work.

